# Modulation of Automatic Alcohol Approach Tendencies using Single-Session 10 Hz rTMS over the Right dLPFC

**DOI:** 10.64898/2026.04.09.717508

**Authors:** Adarsh K. Verma, Adith Deva Kumar, Usha Chivukula, Neeraj Kumar

## Abstract

**Background:** Maladaptive drinking is often sustained by automatic approach tendencies toward alcohol cues that override conscious self-control. While cognitive and behavioral modification techniques show some promise, their effects remain limited, highlighting the need for alternative neuromodulatory strategies. The current study examined the feasibility of a single session of 10 Hz repetitive transcranial magnetic stimulation (rTMS) to the right dorsolateral prefrontal cortex (dLPFC) as a targeted approach to reduce automatic alcohol approach tendencies.

**Method:** Forty-five healthy alcohol-using participants completed an alcohol approach– avoidance task (A-AAT) with concurrent electroencephalographic recording before and after active or sham stimulation. Primary analyses focused on participants with baseline alcohol approach tendencies (n = 35).

**Results:** At baseline, individuals with approach tendencies exhibited attenuated N2 and P3b amplitudes to alcohol relative to non-alcohol cues, indicating reduced cognitive control and attentional mechanisms irrespective of group. Following stimulation, active rTMS selectively facilitated alcohol avoidance responses and enhanced prefrontal N2 amplitudes, suggesting strengthened top-down control and protection against repetition-induced automaticity, which was evident in the sham group.

**Conclusion:** These findings suggest that high-frequency rTMS over the right dLPFC can modulate automatic alcohol-related action tendencies by strengthening neural control mechanisms, supporting its further evaluation as a neuromodulatory adjunct for maladaptive drinking. Baseline motivational profiles may additionally influence rTMS response and warrant consideration when tailoring such approaches.

## Introduction

Harmful alcohol use is characterized by the progressive strengthening of automatic approach tendencies toward alcohol-related cues, driven by the sensitization of appetitive motivational systems through repeated alcohol exposure (Cofresí et al., 2019; Fleming & Bartholow, 2014). Within a dual-process framework, these automatic tendencies operate with limited conscious oversight and increasingly override executive regulatory control, resulting in compulsive alcohol-seeking behavior despite adverse consequences (Lannoy et al., 2014; Lindgren et al., 2019). This imbalance between automatic approach responses and deliberate inhibitory control is reflected neurally in attenuated conflict-monitoring and stimulus-evaluation processes, as indexed by reduced right-frontal N2 and centro-parietal P3 event-related potentials during alcohol-related decision-making (Luijten et al., 2014; Petit et al., 2012). Critically, the robustness of these neurocognitive mechanisms underlies the persistence of relapse vulnerability (R. W. Wiers et al., 2011) and motivates the investigation of interventions that directly target approach–avoidance control.

Several cognitive and behavioral interventions have been developed to modify automatic approach tendencies, such as cognitive bias modification (CBM), which attempts to retrain approach biases through repeated practice in pushing away alcohol cues (R. W. Wiers et al., 2011). Although CBM demonstrates efficacy under controlled laboratory conditions, meta-analytic evidence indicates small effect sizes, and the sustainability of these behavioral changes in clinical settings is limited (Boffo et al., 2019; Pan et al., 2026). Importantly, even after such interventions, neurocognitive deficits often persist (Crowe et al., 2020; Schulte et al., 2014), highlighting the resistance of underlying neurocognitive mechanisms to purely behavioral interventions (Everitt & Robbins, 2016). These observations motivate the investigation of neuromodulation approaches that can directly target the neural circuits governing approach– avoidance conflict.

Transcranial direct current stimulation (tDCS), which modulates cortical excitability by applying electrical current, has shown a modest effect in reducing alcohol craving (Chan et al., 2024). However, attempts to directly modify automatic alcohol approach tendencies using tDCS, either as a standalone intervention (e.g., stimulation of the left dorsolateral prefrontal cortex [dLPFC]) or in combination with cognitive bias modification, have yielded largely null findings (Claus et al., 2019; Schwippel et al., 2022). This discrepancy between tDCS efficacy for craving versus approach tendencies may reflect distinct neural mechanisms, where craving primarily involves dysregulated activity in reward circuits modulated through changes in cortical excitability (Koob & Volkow, 2010), whereas automatic approach tendencies represent habitual stimulus-response associations mediated by cortico-striatal circuits (Everitt & Robbins, 2016; Robbins et al., 2012). The limitation of tDCS in addressing habitual approach behaviors may reflect its primarily subthreshold neuromodulatory effects, which alter cortical excitability without inducing the sustained plasticity necessary to reorganize well-established habitual circuits (Nitsche & Paulus, 2011; Stagg & Nitsche, 2011). Repetitive transcranial magnetic stimulation (rTMS) offers a mechanistically distinct approach that may overcome these limitations. Unlike tDCS, high-frequency rTMS (≥5 Hz) induces suprathreshold neuronal depolarization through repeated magnetic pulses, triggering action potentials and producing long-term potentiation-like plasticity in targeted cortical circuits (Halawa et al., 2021). This capacity to induce synaptic changes and modulate cortical excitability is particularly relevant given that automatic approach tendencies are thought to be supported by strongly consolidated stimulus-response associations in prefrontal and striatal circuits that may require a greater degree of plasticity induction than subthreshold techniques can achieve. The neuroplastic effects of rTMS can persist for weeks to months beyond stimulation (Rossi et al., 2009), suggesting potential for enduring changes in habit-dependent circuits. High-frequency rTMS has demonstrated efficacy in reducing alcohol craving and improving cognitive function in alcohol-dependent individuals (Feng et al., 2022; Mishra et al., 2010), yet its potential for modulating automatic approach tendencies remains unexplored.

The right dLPFC represents a particularly promising target for rTMS stimulation based on converging neuroimaging evidence. Functional MRI studies demonstrate that right dLPFC activity distinguishes approach from avoidance responses to alcohol cues, with significantly stronger activation during avoidance than during approach responses (Ernst et al., 2014), suggesting its role in implementing cognitive control over prepotent approach tendencies. Electrophysiological studies further identify the right dLPFC and centro-parietal regions as key nodes in the neural circuitry mediating approach–avoidance conflict (Ernst et al., 2014; Friedman & Robbins, 2022). Task-related modulation of the N2 and P3b components within this network indexes early conflict detection and subsequent controlled stimulus evaluation, respectively (Cao et al., 2017; Petit et al., 2012). Individuals exhibiting alcohol approach tendencies consistently show attenuated N2 and P3b amplitudes during alcohol-related inhibition tasks, indicating compromised engagement of control mechanisms (Domínguez-Centeno et al., 2018; Ernst et al., 2014). Given the evidence that high-frequency rTMS targeting the right dLPFC enhances frontal–parietal functional connectivity (Kazemi et al.,2020), such stimulation may help augment deficient top-down control processes and reduce automatic alcohol approach tendencies. Building on this neurocognitive framework, the present study tested whether modulating right dLPFC activity via high-frequency rTMS can alter automatic alcohol approach behavior and its associated neural signatures.

Specifically, the present study examined whether high-frequency (10 Hz) rTMS applied to the right dLPFC could reduce automatic alcohol approach tendencies. We hypothesized that individuals exhibiting alcohol approach tendencies would show attenuated right-frontal N2 and centro-parietal P3b amplitudes at baseline relative to individuals with avoidance tendencies, as evident in earlier literature, reflecting weakened neural processes supporting conflict detection and controlled stimulus evaluation (Verma et al., 2025). We further hypothesized that excitatory rTMS would enhance these earlier attenuated markers and reduce behavioral alcohol approach tendencies, indicating strengthened top-down control over automatic responses to alcohol cues. Although baseline motivational tendencies were used to characterize participants, primary hypotheses and analyses focused on individuals with approach tendencies, consistent with the study’s central objective of reducing maladaptive alcohol approach behavior.

At baseline, our results demonstrate attenuated neural responses to alcohol stimuli in individuals with approach tendencies, with behavioral (reduced automatic approach tendencies) and neural enhancements following 10 Hz rTMS. However, its effects were limited to the right frontal region in the approach participants. Our study provides proof-of-concept for excitatory rTMS at the rDLPFC as a viable neuromodulatory approach to augment automatic alcohol approach behavior.

## Method

### Participants

An a priori power analysis was performed using G*Power (Faul et al., 2007) to estimate the required sample size for a mixed-design ANOVA. The model included stimulation condition (active vs. sham) as a between-subjects factor and time (pre-vs. post-rTMS) as a within-subjects factor. Based on an assumed medium effect size (f = 0.25), a significance threshold of α = 0.05, and a desired statistical power of 0.80, the analysis indicated that at least 34 participants were needed to detect a stimulation-related effect.

Since automatic alcohol approach or avoidance tendencies can only be identified following administration of the Alcohol Approach–Avoidance Task (A-AAT) and cannot be determined a priori, a larger sample was recruited. Accordingly, 45 male participants (mean age = 23.3 ± 2.94 years) were enrolled in the study to minimize sex-related confounds, such as gendered drinking practices (Cook et al., 2025), competing response tendencies (Weafer et al., 2010), and interventional effects (Unlu et al., 2023), thereby enhancing internal validity and facilitating mechanistic interpretability of the findings. Eligible participants were regular alcohol users, predominantly right-handed or mixed-right-handed, with no self-reported history of neurological or psychiatric illness, and with normal or corrected-to-normal vision. Eligibility for rTMS was assessed using the Standard Screening Questionnaire for rTMS Candidates (Rossi et al., 2011). Alcohol use severity was assessed using the Alcohol Use Disorder Identification Test (AUDIT), where 31 participants showed low-risk, 9 showed moderate-risk, and 4 showed high-risk drinking patterns. None met diagnostic criteria for alcohol dependence at the time of enrolment. To minimize the acute effects of alcohol, participants were required to abstain from drinking for a minimum of 24 hours prior to the experimental session.

Ethical approval for the study was obtained from the Institutional Ethics Committees of the University of Hyderabad (UH/IEC/2025/267) and the Indian Institute of Technology Hyderabad (IITH/IEC/2025/01/13). All procedures adhered to the principles outlined in the Declaration of Helsinki, and written informed consent was obtained from each participant prior to participation.

Following baseline A-AAT assessment, participants demonstrating alcohol approach tendencies were alternately allocated, based on recruitment sequence, to receive either active rTMS (n = 17) or sham stimulation (n = 18). One participant initially allocated to the active stimulation group was excluded from EEG analyses due to excessive recording artifacts. In addition, nine participants who exhibited avoidance tendencies at baseline were assigned to receive active stimulation and were included as a comparison group to examine post-rTMS changes. Given the smaller sample size, analyses involving the avoidance group were considered exploratory and are reported in the Supplementary Material.

### Experimental Design and Task

A mixed factorial experimental design was employed to examine changes in automatic alcohol approach tendencies following high-frequency (10 Hz) rTMS. The design included time (pre-vs. post-rTMS) as a within-subjects factor and stimulation condition (active vs. sham) as a between-subjects factor. Baseline alcohol approach or avoidance tendencies, identified using the Alcohol Approach–Avoidance Task (A-AAT), constituted a measured grouping variable rather than an experimentally manipulated factor. Behavioral and neurophysiological outcomes were assessed using the A-AAT in conjunction with event-related potentials (ERPs). The primary analyses focused on individuals exhibiting alcohol approach tendencies at baseline, consistent with the study’s objective about rTMS influence on automatic approach tendencies, while avoidance tendencies were examined exploratorily. The A-AAT was administered using PsychToolbox-3 in MATLAB (MathWorks, Inc., Natick, MA, USA) to quantify changes in automatic approach and avoidance responses to alcohol-related cues.

### Alcohol Approach-Avoidance Task (A-AAT)

Automatic tendencies toward alcohol-related stimuli were measured using the Alcohol Approach–Avoidance Task (A-AAT), adapted from established paradigms (Kersbergen et al., 2015; Rinck & Becker, 2007). The task utilized images of alcoholic and non-alcoholic beverages as stimuli, with participants responding via joystick movements (pull or push) to indicate approach and avoidance responses, respectively. To enhance the salience of action tendencies, joystick movements were coupled with dynamic visual feedback: pulling the joystick enlarged the image (zoom-in effect), whereas pushing reduced its size (zoom-out effect), creating a perceptual correspondence between motor action and approach–avoidance behavior.

The task comprised 180 trials organized into three sequential blocks. The two experimental blocks (80 trials each) employed counterbalanced stimulus–response mappings. In the first experimental block, participants were instructed to pull (approach) alcohol-related images and push (avoid) non-alcohol images. This mapping was reversed in the second experimental block, requiring participants to push alcohol images and pull non-alcohol images. Between the two experimental blocks, a washout block consisting of 20 trials was administered. During this washout period, participants performed approach–avoidance responses to neutral stimuli (bells and bulbs), pulling bell images and pushing bulb images. This washout procedure was implemented to eliminate potential carryover effects of the pull/push motor practice from the first block that might influence performance in the second block.

Each trial began with a fixation cross presented centrally for 500 ms, followed by stimulus presentation lasting up to 3000 ms. Participants were required to keep the joystick in a neutral position for 200 ms before stimulus onset to minimize premature responses. During practice trials, participants received immediate visual feedback with a green tick for correct responses or a red cross for incorrect responses. During the experimental trials, feedback was presented only following incorrect responses to promote accuracy in further trials.

### 10 Hz rTMS Protocol

Transcranial magnetic stimulation was administered using the Deymed DuoMAG XT-100 system equipped with a figure-of-eight coil. The stimulation target was the right dorsolateral prefrontal cortex (dLPFC), identified using the BeamF3 method (Beam et al., 2009) adapted for the right hemisphere, and matched with the F4 electrode site according to the 10-20 EEG system. Precise coil positioning was achieved using a neuronavigation system (NDI Polaris Vicra) following registration of each participant’s head to the MNI head template based on anatomical landmarks (nasion–inion and tragus–tragus).

The resting motor threshold (RMT) was determined by delivering single-pulse TMS over the left primary motor cortex (C3 site) to elicit motor-evoked potentials (MEPs) in the contralateral right first dorsal interosseous (FDI) muscle. RMT was defined as the minimum stimulation intensity producing MEPs of at least 50 μV amplitude in five out of ten consecutive pulses. The excitatory high-frequency rTMS protocol consisted of 1,560 pulses delivered across 40 trains of 10 Hz stimulation, with each train lasting 3.9 seconds and an inter-train interval of 26.1 seconds. Stimulation intensity was set at 110% of individual RMT (Vanderhasselt et al., 2007). In the sham stimulation group, the coil was positioned vertically at a 90° angle, contacting the scalp with only one edge.

### EEG Recording and Preprocessing

Electroencephalographic activity was recorded using the Brain Products actiChamp system with 64 active Ag/AgCl electrodes positioned according to the international 10-20 system. Data were acquired at a sampling rate of 1000 Hz with impedances maintained below 10 kΩ throughout recording.

EEG data preprocessing was carried out offline using the EEGLAB toolbox (Delorme & Makeig, 2004) in MATLAB. The preprocessing steps included application of a 50 Hz notch filter to attenuate line noise, independent component analysis (ICA) using the runica algorithm to identify and remove components associated with ocular and muscle artifacts, and segmentation of continuous data into epochs spanning -1000 to 2000 ms relative to stimulus onset. To enhance the signal-to-noise ratio for target ERP components, a bandpass filter of 0.2– 10 Hz was applied (Zhang et al., 2024).

For ERP analysis, regions of interest (ROIs) were defined a priori. The left frontal (LF) ROI, representing the left dorsolateral prefrontal cortex, comprised electrodes F1, F3, F5, F7, AF3, FC3, and FC5. The right frontal (RF) ROI, corresponding to the right dorsolateral prefrontal cortex, included electrodes F2, F4, F6, F8, AF4, FC4, and FC6. The centro-parietal (CP) ROI was calculated as the average signal across electrodes CPz and Pz. Visual inspection of grand-averaged waveforms guided the selection of temporal windows for ERP quantification. The N2 component was measured as the mean amplitude between 200–400 ms post-stimulus at frontal ROIs (LF and RF), while the P3b component was extracted from the CP ROI in the 350–550 ms window.

### Statistical Analysis

Median reaction times (RTs) were computed per participant for each stimulus × action combination to reduce the influence of extreme values. The Alcohol Approach–Avoidance Index (AAI) was derived by subtracting the non-alcohol approach score (pull minus push RT) from the alcohol approach score, with positive values indicating automatic approach tendencies. Preliminary equivalence analyses used one-way ANOVA for age, chi-square or Fisher’s exact tests for categorical variables, and parametric or nonparametric tests for baseline behavioral and neural measures, depending on whether normality and homogeneity assumptions were satisfied. Baseline ERP amplitudes were examined using mixed-effects ANOVAs with stimulus, movement, and group as fixed factors and participant as a random factor.

rTMS-related behavioral and neural effects were analyzed using aligned rank transformation ANOVAs (ART-ANOVA; Wobbrock et al., 2011), with assessment phase (pre vs. post) as a within-subject factor and stimulation group as a between-subject factor. Significant effects were followed by post hoc Wilcoxon signed-rank or t-tests, as appropriate, with effect sizes reported as Wilcoxon r (Tomczak & Tomczak, 2014) or Cohen’s d. P-values were corrected for multiple comparisons using the Benjamini–Hochberg procedure. Bayes factors (BF10; Morey & Rouder, 2012) were computed throughout to complement frequentist inference, particularly for non-significant effects. All analyses were conducted in R (Version 4.3.1; R Core Team, 2024).

## Results

Automatic alcohol approach tendencies are a key feature of maladaptive drinking behavior. Recent evidence points to the right dLPFC in modulating these automatic tendencies (C. E. Wiers et al., 2014; Xia et al., 2021). The current study explores the utility of 10 Hz rTMS at the right dLPFC to reduce automatic approach tendencies towards alcohol cues in healthy alcohol-using individuals by enhancing top-down cognitive control and reducing automaticity in alcohol approach behavior.

### Participant Characteristics and Group Allocation

The study recruited 45 participants, of whom 9 demonstrated avoidance tendencies during the initial A-AAT administration. These participants received 10 Hz stimulation to explore tendency-specific effects, and their results are presented in the Supplementary Material due to the limited sample size. The remaining 36 participants, who exhibited approach tendencies toward alcohol cues, were alternately allocated to either active (10 Hz rTMS) or sham stimulation groups. One participant’s data from the active stimulation group was excluded due to substantial artifacts in the electroencephalography recording, yielding a final sample of 17 participants in the active stimulation group and 18 in the sham group.

Preliminary analyses confirmed no significant differences between the three groups (active, sham, and avoidance) in key demographic variables, i.e., age (F(2, 41) = 2.03, p = 0.145, BF10 = 0.716), AUDIT risk pattern (Fisher’s p = 0.77; χ^2^(4, N = 43) = 2.63, p = 0.62), and education level (Fisher’s p = 0.41; χ^2^(4, N = 43) = 4.22, p = 0.38). Descriptive statistics for each group are presented in Table 1, demonstrating comparable baseline characteristics across conditions.

**Table 1.** Demographics and baseline characteristics of participants across groups.

|  | Approach-Active | Approach-Sham | Avoidance-Active |
| --- | --- | --- | --- |
| N [% of total] | 17 [38.6] | 18 [40.9] | 9 [20.4] |
| Age (mean ± SD) | 23.1 ± 2.41 | 24.4 ± 2.12 | 22.9 ± 2.67 |
| AAI (Mean ± SE) | 0.115 ± 0.015 | 0.128 ± 0.033 | -0.057 ± 0.021 |
| AUDIT Score (Mean ± SE) | 6.94 ± 1.47 | 8.17 ± 1.75 | 6.11 ± 1.27 |
| AUDIT Classification (n [% within group]) |  |  |  |
| low risk (1-7) | 13 [76.5] | 11 [61.1] | 7 [77.78] |
| moderate risk (8-14) | 3 [17.6] | 4 [22.2] | 2 [22.22] |
| high risk (15+) | 1 [5.8] | 3 [16.7] | - |
| Educational program (n [% within group]) |  |  |  |
| Undergraduate | 10 [58.8] | 5 [27.8] | 5 [55.6] |
| Postgraduate | 5 [29.4] | 8 [44.4] | 3 [33.3] |
| Doctorate | 2 [11.8] | 5 [27.8] | 1 [11.1] |

*Note.* AAI = Alcohol Approach-Avoidance Index; AUDIT = Alcohol Use Disorder Identification Test.

### Group Equivalence in Behavioral and Neural Responses at Baseline

Baseline behavioral responses (AAI) did not differ significantly between active and sham groups (median difference = 0.046, W = 179, p = 0.242, BF10 = 0.363), confirming comparable automatic approach tendencies toward alcohol cues prior to rTMS. Baseline neural responses similarly showed no group differences across ERPs, i.e., LF N2 amplitudes (F(1, 33) = 0.484, p = 0.491, BF10 = 0.402), RF N2 amplitudes (F(1, 33) = 1.697, p = 0.202, BF10 = 1.095), and CP P3b amplitudes (F(1, 33) = 0.973, p = 0.331, BF10 = 0.373).

### Alcohol-Specific RF N2 and CP P3 Attenuation

Participants in both groups showed robust stimulus-specific N2 responses. At the RF region, a significant main effect of stimulus emerged (F(1, 33) = 41.435, p < 0.001), with lower N2 amplitudes for alcohol versus non-alcohol cues (median difference = 3.820, W = 584, p < 0.001, r = 0.745), suggesting diminished controlled processing for alcohol stimuli. No such effect was observed at LF (F(1, 33) = 0.912, p = 0.347, BF10 = 0.196), and no three-way group × stimulus × action interaction emerged at either LF (F(1, 33) = 0.004, p = 0.952, BF10 = 0.563) or RF (F(1, 33) = 1.546, p = 0.222, BF10 = 0.402).

At the CP region, P3b amplitudes were significantly lower for alcohol than non-alcohol cues (F(1, 33) = 19.551, p < 0.001, ω^2^ = 0.08; median difference = −4.09, W = 83, p < 0.001, r = 0.642), potentially reflecting the automaticity in alcohol-cue processing, with lower allocation of attentional resources. Although a main effect of movement also emerged (F(1, 33) = 7.613, p = 0.009), meaningful follow-up within-group analyses revealed no significant pull–push differences for either alcohol (median difference = 3.10; W = 114, p = 0.080, BF10 = 1.013) or non-alcohol cues (median difference = 2.59; W = 107, p = 0.156, BF10 = 0.859) in the active group or the sham group (alcohol: median difference = 0.895; W = 87, p = 0.597, BF10 = 0.282; non-alcohol: median difference = 3.460; W = 115, p = 0.207, BF10 = 0.597). No three-way interaction was observed (F(1, 33) = 0.004, p = 0.948, BF10 = 0.311), indicating the omnibus movement effect reflects variance across the full factorial design rather than systematic pull–push differences within theoretically relevant conditions.

### Behavioral Effects of 10 Hz rTMS: Reduction in Automatic Approach Tendencies by Strengthening Avoidance

To assess rTMS efficacy at the behavioral level, changes in the Alcohol Approach Index (AAI) were examined across groups (active and sham) x assessment phase (pre- and post-rTMS). A significant group × assessment phase interaction (F(1, 32) = 8.286, p = 0.007) revealed differential behavioral trajectories between groups. Participants receiving active 10 Hz rTMS showed a significant reduction in automatic approach tendencies toward alcohol (paired median difference = −0.022, W = 30, p = 0.027, r = 0.405), whereas the sham group showed no such change (paired median difference = 0.009, W = 99, p = 0.306, BF10 = 0.368; Figure 1).

**Figure 1.**
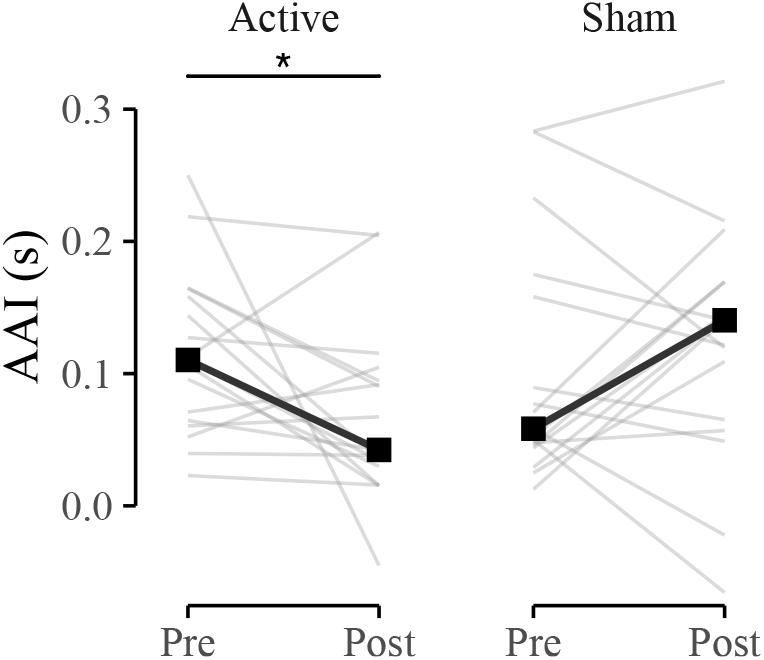
Active 10 Hz rTMS reduces alcohol-approach tendencies in participants with baseline approach bias. Individual AAI scores (lighter lines) and group medians (darker lines) are shown pre- and post-intervention.

To understand the mechanism underlying this AAI reduction, effects were decomposed at the level of individual trial types comprising the index. A significant three-way interaction between trial type (alcohol pull, alcohol push, non-alcohol pull, and non-alcohol push), assessment phase (pre- and post-rTMS), and group (active and sham) (F(3, 224) = 28.348, p < 0.001) indicated that rTMS effects varied systematically across conditions. In the active group, the AAI reduction was driven specifically by faster alcohol push responses post-rTMS compared to the baseline (paired median difference = −0.038, W = 29, p = 0.037, r = 0.545), with no significant change in alcohol pull responses (paired median difference = −0.005, W = 95, p = 0.404, BF10 = 0.406; Figure 2 Panel A). No significant changes were observed for non-alcohol stimuli, for either push (median difference = -0.020, W = 38, p = 0.082, BF10 = 1.04) or pull trials (median difference = -0.029, W = 37, p = 0.082, BF10 = 1.46; Figure 2 Panels B). This suggests that rTMS selectively regulated the capacity to execute alcohol avoidance actions, rather than producing a generic effect on all actions and stimulus types.

**Figure 2.**
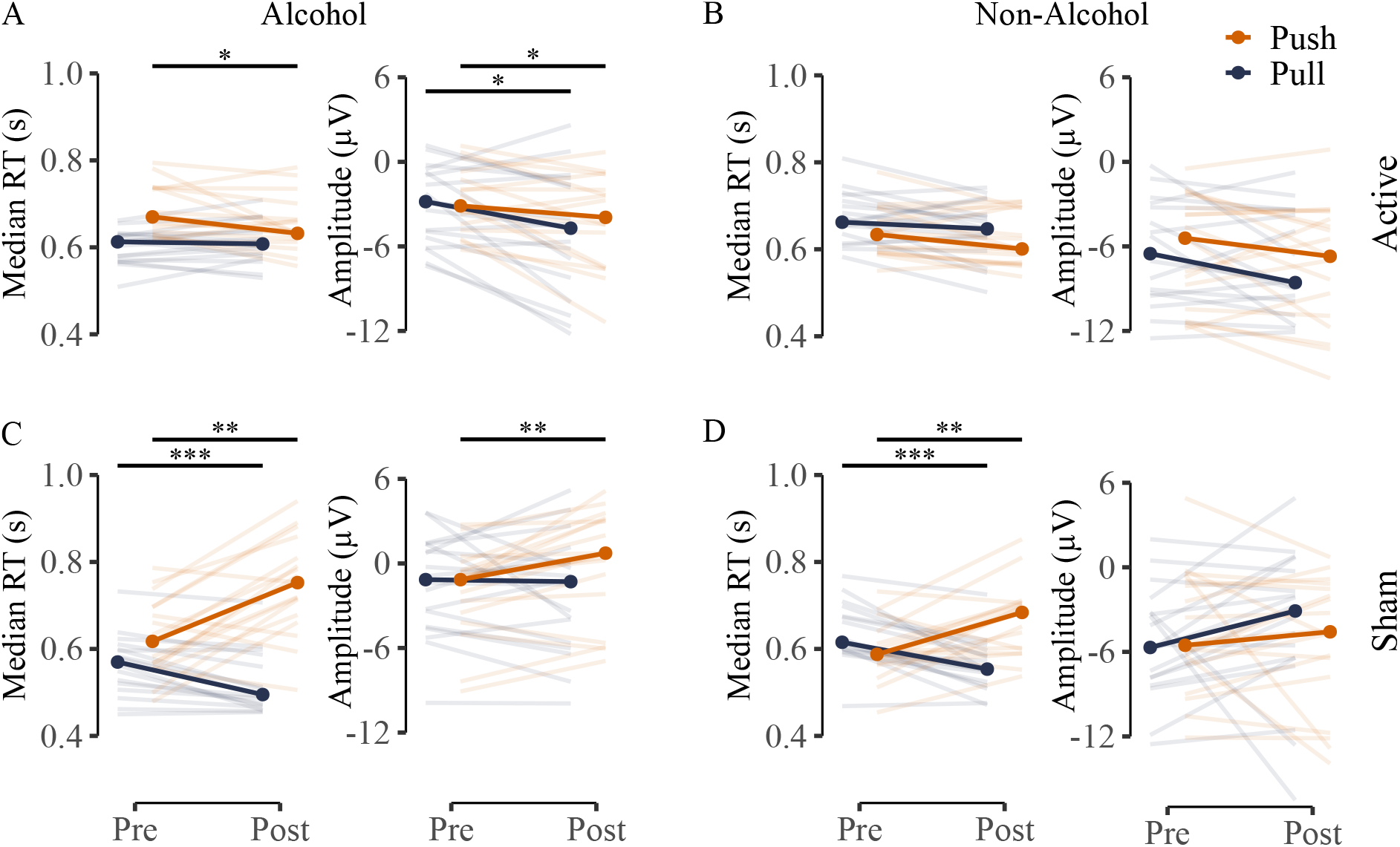
In the active stimulation group, the reduction in AAI was primarily driven by enhanced alcohol avoidance, reflected in faster push responses (reduced RTs) and enhanced RF N2 amplitudes for alcohol trials (Panel A). No comparable changes were observed for non-alcohol stimuli (Panel B). In contrast, the sham group showed a general facilitation of pull responses and relative slowing of push responses (Panels C and D). Lighter lines represent individual participants, and darker lines indicate group medians.

### Repetition Effects Reflected in Sham Group

The sham group exhibited a pattern consistent with task repetition effects, where repeated exposure without active neuromodulation led to a response time trade-off. Pull responses became faster for both alcohol (paired median difference = −0.075, W = 4, p < 0.001, r = 0.832) and non-alcohol stimuli (paired median difference = −0.062, W = 1, p < 0.001, r = 0.867), while push responses slowed for both alcohol (paired median difference = 0.135, W = 143, p = 0.002, r = 0.763) and non-alcohol stimuli (paired median difference = 0.096, W = 136, p = 0.006, r = 0.683), favoring appetitive alcohol approach over avoidance (Figure 2, Panels C and D). Direct post-rTMS between-group comparisons confirmed these divergent effects, where active participants showed slower alcohol pull (paired median difference = 0.113, W = 250, p = 0.001, r = 0.623) and faster alcohol push responses than sham participants (paired median difference = −0.120, W = 66, p = 0.016, r = 0.464). These results indicate that 10 Hz rTMS to the right dLPFC selectively enhanced alcohol avoidance responding, while the sham condition produced the opposite pattern, i.e., strengthened automatic approach and weakened avoidance, consistent with repetition-driven automaticity.

### Neural Effects of 10 Hz rTMS: Enhanced Cognitive Control During Alcohol-Cue Processing

To examine the neural mechanisms underlying behavioral changes, we investigated whether rTMS modulated cognitive control processes reflected by N2 amplitude at frontal regions and attentional allocation indexed by P3b amplitude at the centroparietal region. At the centroparietal region, neither a main effect of rTMS (F(1, 231) = 0.182, p = 0.670, BF10 = 0.138) nor a four-way interaction between assessment phase, group, stimulus type, and action (F(1, 231) = 0.130, p = 0.719, BF10 = 0.464) reached significance, indicating that P3b-indexed attentional allocation was not systematically modulated by rTMS. At the RF region, however, a significant four-way interaction emerged (F(1, 231) = 4.051, p = 0.045), indicating that rTMS specifically modulated prefrontal cognitive control during alcohol-related approach and avoidance.

Within-group comparisons revealed different neural trajectories across active and sham groups. In the sham group, N2 amplitude during alcohol push trials was significantly reduced post-rTMS relative to baseline (paired median difference = 1.88, W = 11, p = 0.003, r = 0.765), Figure 2 Panel C. This reduction suggests diminished cognitive control during alcohol avoidance and aligns with the corresponding behavioral slowing of push responses. In contrast, the active group showed enhanced N2 amplitudes for both alcohol pull (paired median difference = −1.89, W = 127, p = 0.06, BF10= 4.78, r = 0.580) and push actions (paired median difference = −0.816, W = 122, p = 0.081, BF10 = 3.21, r = 0.522), Figure 2 Panel A. Between-group comparisons post-rTMS confirmed enhanced N2 amplitudes in the active relative to the sham group during both alcohol pull (paired median difference = 3.42, W = 90, p = 0.101, BF10 = 1.60, r = 0.351) and push trials (paired median difference = 4.67, W = 57, p = 0.008, r = 0.536). This active-sham group difference was primarily driven by the attenuation of N2 responses in the sham group, rather than a uniform enhancement in the active group. This pattern suggests that without subsequent neural intervention, repeated exposure to alcohol cues may reduce prefrontal control while increasing behavioral automaticity, where neural enhancements through 10 Hz rTMS can lead to facilitation of alcohol push (avoidance) responses.

## Discussion

Automatic approach tendencies toward alcohol cues represent a core, yet intervention-resistant, mechanism in problematic drinking (Boffo et al., 2019). The present study demonstrates that, in healthy alcohol-using individuals, a single session of 10 Hz rTMS targeting the right dLPFC may meaningfully disrupt these tendencies, not by suppressing automatic approach directly but likely through rDLPFC-related strengthening of avoidance responses to alcohol cues and prefrontal cognitive control. Crucially, this effect becomes particularly meaningful when considered in contrast to the sham condition, where repeated task exposure without stimulation produced the opposite effect, i.e., accelerated approach and deteriorating avoidance, mirroring early-stage habit formation (Buabang et al., 2025; Yamada & Toda, 2023). The divergence underscores both the therapeutic utility of excitatory prefrontal stimulation and the potential risk of unguided cue exposure in clinical settings, and positions right dLPFC rTMS as a promising add-on while targeting alcohol-related automaticity.

### Neurophysiological Signatures of Automatic Alcohol Approach Tendencies

The baseline attenuation of right-frontal N2 and centro-parietal P3b amplitudes to alcohol cues in individuals with approach tendencies aligns with dual-process models proposing that automatic approach behaviors reflect weakened top-down cognitive control over prepotent responses (Lindgren et al., 2019). The N2 component, generated primarily in the anterior cingulate cortex and prefrontal regions, is thought to reflect conflict detection and the recruitment of cognitive control when response competition occurs (Carter & van Veen, 2007; Folstein & Van Petten, 2008). The P3b component, with its centro-parietal distribution, reflects the controlled evaluation of stimulus significance and allocation of attentional resources (Polich, 2007). The N2 and P3b attenuation combinedly suggests that alcohol cues might have been processed through relatively automatic, stimulus-driven pathways that bypass deliberate evaluation, facilitating rapid approach responses without adequate regulatory engagement. The specificity of these neural alterations to alcohol stimuli, with no comparable effects for non-alcohol cues, indicates that approach tendencies involve cue-specific disruptions in cognitive control rather than generalized executive dysfunction, consistent with dual-process accounts in which substance-related cues acquire privileged access to behavioral output systems through repeated pairing with reward (Lindgren et al., 2019; McClure & Bickel, 2014).

While cognitive control is typically considered a bilateral process, converging evidence implicates the right prefrontal cortex, especially in avoidance regulation and inhibitory control (Friedman & Robbins, 2022; Watson et al., 2018), and the right dLPFC has been specifically associated with successful avoidance of alcohol cues in neuroimaging studies (Ernst et al., 2014). Our baseline findings extend this work by demonstrating that individuals with approach tendencies show reduced engagement of right frontal control mechanisms when processing alcohol cues, which may explain why the right dLPFC emerged as an effective neuromodulation target in the present study.

### Mechanisms of rTMS-Induced Reduction in Approach Tendencies

Following active rTMS, enhanced right frontal N2 amplitudes for both alcohol pull and push actions indicate that stimulation might have enhanced right dLPFC excitability, leading to increased top-down control recruitment during alcohol cue processing. This finding is consistent with the known role of the right dLPFC in implementing control over habitual responses, particularly in contexts requiring response inhibition or conflict resolution against prepotent responses (Friedman & Robbins, 2022; Watson et al., 2018). The behavioral specificity of rTMS effects, i.e., enhanced alcohol avoidance without significant changes in alcohol approach responses, suggests that enhanced prefrontal control selectively supported the execution of avoidance actions. The parallel N2 enhancement during both pull and push actions toward alcohol, despite behavioral changes being confined to push responses, may indicate that rTMS increased general control over alcohol cues. The behavioral expression of this enhanced control was most apparent where regulatory demand was highest, i.e., during active avoidance of appetitive stimuli.

The absence of centro-parietal P3b modulation following rTMS may reflect the anatomical specificity of single-session rTMS effects, which are typically most pronounced at the stimulation site with weaker propagation to distal cortical regions (Rossi et al., 2009). The centro-parietal P3b is thought to reflect evaluative processing that may require multi-session protocols to produce cumulative effects across functionally connected networks. Alternatively, the sustained P3b attenuation may reflect the continuation of relatively automatic processing for alcohol cues, with enhanced prefrontal control enabling behavioral regulation despite persistent automaticity in later evaluative stages.

### Repetition Effects and the Protective Role of rTMS

The sham group’s pattern of a systematic shift toward faster approach and slower avoidance responses across the sessions highlights an underappreciated risk in approach-avoidance paradigms where repeated alcohol cue exposure without concurrent intervention may inadvertently reinforce the very stimulus-response associations targeted for modification. The appetitive nature of alcohol cues naturally primes approach responses. When these cues are repeatedly paired with pull actions during paradigms such as A-AAT, the existing approach associations appear to be further consolidated, while avoidance of the same appetitive cues becomes increasingly effortful and slow. This pattern aligns with theoretical accounts of habit formation in which repeated stimulus-response pairings lead to progressive automatization and reduced dependence on deliberative control (Dahm et al., 2023; Doñamayor et al., 2022), and raises important questions about whether bias modification tasks risk producing similar reinforcement of approach tendencies over repeated administrations.

Additionally, active rTMS seems to prevent this strengthening of approach responses while simultaneously supporting avoidance facilitation, suggesting that the stimulation might have protected against maladaptive plasticity that would otherwise occur through task repetition. This interpretation is consistent with evidence that rTMS can modulate synaptic plasticity mechanisms, including long-term potentiation in prefrontal circuits, which may have shifted the balance of plasticity toward a more regulatory than automatic approach pathways.

### Theoretical and Clinical Implications

Our findings support dual-process models of addiction, which propose that problematic alcohol use arises from an imbalance between strong automatic approach tendencies and weakened executive control (Lindgren et al., 2019). The reduced right-frontal N2 amplitudes observed at baseline reflect this imbalance, indicating that individuals with elevated approach tendencies engage less prefrontal conflict monitoring when encountering alcohol cues. Further, our results highlight the importance of distinguishing between reducing approach and enhancing avoidance as distinct therapeutic mechanisms. The fact that rTMS enhanced avoidance without significantly reducing approach responses indicates that strengthening active avoidance may be a more tractable target for neuromodulation than suppressing automatic approach. This insight aligns with recent reconceptualizations of addiction treatment, which emphasize building adaptive regulatory capacities rather than simply suppressing maladaptive behaviors (Copersino, 2017; Kwako et al., 2016).

The clinical implications follow directly from these observations. Automatic alcohol cue reactivity has proven resistant to cognitive bias modification, which shows modest effect sizes and limited durability in clinical settings (Boffo et al., 2019). The fact that a single-session rTMS produced measurable behavioral and neural changes suggests it could serve as a useful adjunct to existing treatments. Additionally, given that rTMS appeared to enhance prefrontal excitability and support avoidance consolidation, it may be most effective when delivered prior to structured avoidance practice, potentially amplifying the benefits of cognitive bias modification. Beyond acute application, the sham group’s progressive strengthening of approach responses through repeated unmodulated cue exposure raises an important consideration for relapse prevention. Individuals with alcohol use disorder encounter alcohol cues routinely, and each encounter may reinforce existing approach associations. Periodic rTMS sessions might help maintain prefrontal control capacity and prevent the gradual automatization of approach responses that occurs through repeated cue exposure. This preventive application would represent a novel use of neuromodulatory approaches, focusing on maintaining adaptive neural states that resist automatic habit formation rather than symptom reduction.

### Limitations and Future Directions

Several limitations should be considered when interpreting our findings. First, our sample consisted of healthy non-dependent alcohol users. Alcohol-dependent individuals may have more entrenched habitual approach behaviors supported by more extensive neuroadaptations in prefrontal and striatal circuits that may require multiple rTMS sessions, higher stimulation intensities, or different stimulation targets to achieve comparable effects. Additionally, our sample consisted exclusively of male participants, limiting generalizability to female individuals. Future studies should examine rTMS effects in female participants.

Another limitation is that the present study assessed only immediate post-rTMS effects, precluding conclusions about the durability of behavioral and neural changes. Longitudinal studies tracking approach tendencies and neural responses over days, weeks, and months following rTMS are essential for determining optimal treatment protocols and understanding the time course of therapeutic effects. Finally, our study focused on a single neuromodulation protocol (10 Hz rTMS to right dLPFC), leaving open questions about other brain sites and protocols such as theta burst stimulations.

## Conclusion

This study provides evidence that high-frequency rTMS applied to the right dorsolateral prefrontal cortex can reduce automatic alcohol approach tendencies with corresponding neurophysiological enhancements. Individuals with approach tendencies showed compromised neural processing of alcohol cues at baseline, characterized by attenuated right frontal N2 and centro-parietal P3b amplitudes. A single session of 10 Hz rTMS enhanced neural responses in the right frontal region and reduced automatic approach tendencies toward alcohol, primarily by facilitating avoidance responses rather than suppressing approach. Critically, rTMS also protected against detrimental repetition effects observed in the sham group, wherein repeated task exposure strengthened automatic approach responses. From a clinical perspective, these results suggest excitatory rTMS as a potential intervention for addressing approach tendencies by integration with behavioral interventions to enhance neuroplasticity during cognitive retraining. Future research should examine effects in alcohol-dependent populations, explore optimal integration with behavioral treatments, and determine the longevity of single-session versus multi-session protocols.

## Acknowledgment

This work was supported by the Ministry of Education, Government of India, through the Prime Minister’s Research Fellowship (PMRF) awarded to AKV.

## References

Beam, W., Borckardt, J. J., Reeves, S. T., & George, M. S. (2009). An efficient and accurate new method for locating the F3 position for prefrontal TMS applications. Brain Stimulation, 2(1), 50–54. 10.1016/j.brs.2008.09.006

Boffo, M., Zerhouni, O., Gronau, Q. F., van Beek, R. J. J., Nikolaou, K., Marsman, M., & Wiers, R. W. (2019). Cognitive Bias Modification for Behavior Change in Alcohol and Smoking Addiction: Bayesian Meta-Analysis of Individual Participant Data. Neuropsychology Review, 29(1), 52–78. 10.1007/s11065-018-9386-4

Buabang, E. K., Donegan, K. R., Rafei, P., & Gillan, C. M. (2025). Leveraging cognitive neuroscience for making and breaking real-world habits. Trends in Cognitive Sciences, 29(1), 41–59. 10.1016/j.tics.2024.10.006

Cao, Y., Cao, X., Yue, Z., & Wang, L. (2017). Temporal and spectral dynamics underlying cognitive control modulated by task-irrelevant stimulus-response learning. Cognitive, Affective, & Behavioral Neuroscience, 17(1), 158–173. 10.3758/s13415-016-0469-5

Carter, C. S., & van Veen, V. (2007). Anterior cingulate cortex and conflict detection: An update of theory and data. Cognitive, Affective, & Behavioral Neuroscience, 7(4), 367–379. 10.3758/CABN.7.4.367

Chan, Y.-H., Chang, H.-M., Lu, M.-L., & Goh, K. K. (2024). Targeting cravings in substance addiction with transcranial direct current stimulation: insights from a meta-analysis of sham-controlled trials. Psychiatry Research, 331, 115621. 10.1016/j.psychres.2023.115621

Claus, E. D., Klimaj, S. D., Chavez, R., Martinez, A. D., & Clark, V. P. (2019). A Randomized Trial of Combined tDCS Over Right Inferior Frontal Cortex and Cognitive Bias Modification: Null Effects on Drinking and Alcohol Approach Bias. Alcoholism: Clinical and Experimental Research, 43(7), 1591–1599. 10.1111/acer.14111

Cofresí, R. U., Bartholow, B. D., & Piasecki, T. M. (2019). Evidence for incentive salience sensitization as a pathway to alcohol use disorder. Neuroscience & Biobehavioral Reviews, 107, 897–926. 10.1016/j.neubiorev.2019.10.009

Cook, M., Pennay, A., Caluzzi, G., Cooklin, A., MacLean, S., Riordan, B., Torney, A., & Callinan, S. (2025). Examining gender in alcohol research: A systematic review of gender differences in how men and women are studied in alcohol research. International Journal of Drug Policy, 138, 104763. 10.1016/j.drugpo.2025.104763

Copersino, M. L. (2017). Cognitive mechanisms and therapeutic targets of addiction. Current Opinion in Behavioral Sciences, 13, 91–98. 10.1016/j.cobeha.2016.11.005

Crowe, S. F., Cammisuli, D. M., & Stranks, E. K. (2020). Widespread Cognitive Deficits in Alcoholism Persistent Following Prolonged Abstinence: An Updated Meta-analysis of Studies That Used Standardised Neuropsychological Assessment Tools. Archives of Clinical Neuropsychology, 35(1), 31–45. 10.1093/arclin/acy106

Dahm, S. F., Hyna, H., & Krause, D. (2023). Imagine to automatize: automatization of stimulus–response coupling after action imagery practice in implicit sequence learning. Psychological Research, 87(7), 2259–2274. 10.1007/s00426-023-01797-w

Delorme, A., & Makeig, S. (2004). EEGLAB: an open source toolbox for analysis of single-trial EEG dynamics including independent component analysis. Journal of Neuroscience Methods, 134(1), 9–21. 10.1016/j.jneumeth.2003.10.009

Domínguez-Centeno, I., Jurado-Barba, R., Sion, A., Martinez-Maldonado, A., Castillo-Parra, G., López-Muñoz, F., Rubio, G., & Martinez-Gras, I. (2018). P3 Component as a Potential Endophenotype for Control Inhibition in Offspring of Alcoholics. Alcohol and Alcoholism, 53(6), 699–706. 10.1093/alcalc/agy051

Doñamayor, N., Ebrahimi, C., Arndt, V. A., Weiss, F., Schlagenhauf, F., & Endrass, T. (2022). Goal-Directed and Habitual Control in Human Substance Use: State of the Art and Future Directions. Neuropsychobiology, 81(5), 403–417. 10.1159/000527663

Ernst, L. H., Plichta, M. M., Dresler, T., Zesewitz, A. K., Tupak, S. V., Haeussinger, F. B., Fischer, M., Polak, T., Fallgatter, A. J., & Ehlis, A. (2014). Prefrontal correlates of approach preferences for alcohol stimuli in alcohol dependence. Addiction Biology, 19(3), 497–508. 10.1111/adb.12005

Everitt, B. J., & Robbins, T. W. (2016). Drug Addiction: Updating Actions to Habits to Compulsions Ten Years On. Annual Review of Psychology, 67(1), 23–50. 10.1146/annurev-psych-122414-033457

Faul, F., Erdfelder, E., Lang, A. G., & Buchner, A. (2007). G*Power 3: A flexible statistical power analysis program for the social, behavioral, and biomedical sciences. Behavior Research Methods, 39(2), 175–191. 10.3758/BF03193146

Feng, Z., Wu, Q., Wu, L., Zeng, T., Yuan, J., Wang, X., Kang, C., & Yang, J. (2022). Effect of High-Frequency Repetitive Transcranial Magnetic Stimulation on Visual Selective Attention in Male Patients With Alcohol Use Disorder After the Acute Withdrawal. Frontiers in Psychiatry, 13. 10.3389/fpsyt.2022.869014

Fleming, K. A., & Bartholow, B. D. (2014). Alcohol cues, approach bias, and inhibitory control: Applying a dual process model of addiction to alcohol sensitivity. Psychology of Addictive Behaviors, 28(1), 85–96. 10.1037/a0031565

Folstein, J. R., & Van Petten, C. (2008). Influence of cognitive control and mismatch on the N2 component of the ERP: A review. Psychophysiology, 45(1), 152–170. 10.1111/j.1469-8986.2007.00602.x

Friedman, N. P., & Robbins, T. W. (2022). The role of prefrontal cortex in cognitive control and executive function. Neuropsychopharmacology, 47(1), 72–89. 10.1038/s41386-021-01132-0

Halawa, I., Reichert, K., Aberra, A. S., Sommer, M., Peterchev, A. V., & Paulus, W. (2021). Effect of Pulse Duration and Direction on Plasticity Induced by 5 Hz Repetitive Transcranial Magnetic Stimulation in Correlation With Neuronal Depolarization. Frontiers in Neuroscience, 15. 10.3389/fnins.2021.773792

Kazemi, R., Rostami, R., Dehghan, S., Nasiri, Z., Lotfollahzadeh, S. L., Hadipour, A., Khomami, S., Ishii, R., & Ikeda, S. (2020). Alpha frequency rTMS modulates theta lagged nonlinear connectivity in dorsal attention network. Brain Research Bulletin, 162, 271–281. 10.1016/j.brainresbull.2020.06.018

Kersbergen, I., Woud, M. L., & Field, M. (2015). The validity of different measures of automatic alcohol action tendencies. Psychology of Addictive Behaviors, 29(1), 225–230. 10.1037/adb0000009

Koob, G. F., & Volkow, N. D. (2010). Neurocircuitry of Addiction. Neuropsychopharmacology, 35(1), 217–238. 10.1038/npp.2009.110

Kwako, L. E., Momenan, R., Litten, R. Z., Koob, G. F., & Goldman, D. (2016). Addictions Neuroclinical Assessment: A Neuroscience-Based Framework for Addictive Disorders. Biological Psychiatry, 80(3), 179–189. 10.1016/j.biopsych.2015.10.024

Lannoy, S., Billieux, J., & Maurage, P. (2014). Beyond Inhibition: A Dual-Process Perspective to Renew the Exploration of Binge Drinking. Frontiers in Human Neuroscience, 8. 10.3389/fnhum.2014.00405

Lindgren, K. P., Hendershot, C. S., Ramirez, J. J., Bernat, E., Rangel-Gomez, M., Peterson, K. P., & Murphy, J. G. (2019). A dual process perspective on advances in cognitive science and alcohol use disorder. Clinical Psychology Review, 69, 83–96. 10.1016/j.cpr.2018.04.002

Luijten, M., Machielsen, M., Veltman, D., Hester, R., de Haan, L., & Franken, I. (2014). Systematic review of ERP and fMRI studies investigating inhibitory control and error processing in people. Journal of Psychiatry & Neuroscience, 39(3), 149–169. 10.1503/jpn.130052

McClure, S. M., & Bickel, W. K. (2014). A dual-systems perspective on addiction: contributions from neuroimaging and cognitive training. Annals of the New York Academy of Sciences, 1327(1), 62–78. 10.1111/nyas.12561

Mishra, B. R., Nizamie, S. H., Das, B., & Praharaj, S. K. (2010). Efficacy of repetitive transcranial magnetic stimulation in alcohol dependence: a sham-controlled study. Addiction, 105(1), 49–55. 10.1111/j.1360-0443.2009.02777.x

Morey, R. D., & Rouder, J. N. (2012). BayesFactor: Computation of Bayes Factors for Common Designs. 10.32614/CRAN.package.BayesFactor

Nitsche, M. A., & Paulus, W. (2011). Transcranial direct current stimulation – update 2011. Restorative Neurology and Neuroscience, 29(6), 463–492. 10.3233/RNN-2011-0618

Pan, T., Zhao, X., Bartoš, F., Larsen, H., Manning, V., Boffo, M., & Wiers, R. W. (2026). Cognitive bias modification in alcohol use disorder and problematic drinking: A revised and updated IPD Bayesian meta-analysis. Clinical Psychology Review, 124, 102709. 10.1016/j.cpr.2026.102709

Petit, G., Kornreich, C., Noël, X., Verbanck, P., & Campanella, S. (2012). Alcohol-Related Context Modulates Performance of Social Drinkers in a Visual Go/No-Go Task: A Preliminary Assessment of Event-Related Potentials. PLoS ONE, 7(5), e37466. 10.1371/journal.pone.0037466

Polich, J. (2007). Updating P300: An integrative theory of P3a and P3b. Clinical Neurophysiology, 118(10), 2128–2148. 10.1016/j.clinph.2007.04.019

R Core Team. (2024). R: A language and environment for statistical computing. R Foundation for Statistical Computing. https://www.r-project.org/

Rinck, M., & Becker, E. S. (2007). Approach and avoidance in fear of spiders. Journal of Behavior Therapy and Experimental Psychiatry, 38(2), 105–120. 10.1016/J.JBTEP.2006.10.001

Robbins, T. W., Gillan, C. M., Smith, D. G., de Wit, S., & Ersche, K. D. (2012). Neurocognitive endophenotypes of impulsivity and compulsivity: towards dimensional psychiatry. Trends in Cognitive Sciences, 16(1), 81–91. 10.1016/j.tics.2011.11.009

Rossi, S., Hallett, M., Rossini, P. M., & Pascual-Leone, A. (2009). Safety, ethical considerations, and application guidelines for the use of transcranial magnetic stimulation in clinical practice and research. Clinical Neurophysiology, 120(12), 2008–2039. 10.1016/j.clinph.2009.08.016

Rossi, S., Hallett, M., Rossini, P. M., & Pascual-Leone, A. (2011). Screening questionnaire before TMS: An update. Clinical Neurophysiology, 122(8), 1686. 10.1016/j.clinph.2010.12.037

Schulte, M. H. J., Cousijn, J., den Uyl, T. E., Goudriaan, A. E., van den Brink, W., Veltman, D. J., Schilt, T., & Wiers, R. W. (2014). Recovery of neurocognitive functions following sustained abstinence after substance dependence and implications for treatment. Clinical Psychology Review, 34(7), 531–550. 10.1016/j.cpr.2014.08.002

Schwippel, T., Schroeder, P. A., Hasan, A., & Plewnia, C. (2022). Implicit measures of alcohol approach and drinking identity in alcohol use disorder: A preregistered double-blind randomized trial with cathodal transcranial direct current stimulation (tDCS). Addiction Biology, 27(4). 10.1111/adb.13180

Stagg, C. J., & Nitsche, M. A. (2011). Physiological Basis of Transcranial Direct Current Stimulation. The Neuroscientist, 17(1), 37–53. 10.1177/1073858410386614

Tomczak, E., & Tomczak, M. (2014). The need to report effect size estimates revisited. An overview of some recommended measures of effect size. TRENDS in Sport Sciences, 21(1), 19–25. https://tss.awf.poznan.pl/The-need-to-report-effect-size-estimates-revisited-An-overview-of-some-recommended,188960,0,2.html

Unlu, H., Macaron, M. M., Ayraler Taner, H., Kaba, D., Akin Sari, B., Schneekloth, T. D., Leggio, L., & Abulseoud, O. A. (2023). Sex difference in alcohol withdrawal syndrome: a scoping review of clinical studies. Frontiers in Psychiatry, 14. 10.3389/fpsyt.2023.1266424

Vanderhasselt, M.-A., De Raedt, R., Baeken, C., Leyman, L., Clerinx, P., & D’haenen, H. (2007). The influence of rTMS over the right dorsolateral prefrontal cortex on top-down attentional processes. Brain Research, 1137, 111–116. 10.1016/j.brainres.2006.12.050

Verma, A. K., Kumar, A. D., Chivukula, U., & Kumar, N. (2025). Neural mechanisms underlying approach and avoidance tendencies in alcohol use among males: An electrophysiological investigation. Biological Psychology, 202, 109144. 10.1016/j.biopsycho.2025.109144

Watson, P., van Wingen, G., & de Wit, S. (2018). Conflicted between Goal-Directed and Habitual Control, an fMRI Investigation. Eneuro, 5(4), ENEURO.0240-18.2018. 10.1523/ENEURO.0240-18.2018

Weafer, J. A., Miller, M., & T. Fillmore, M. (2010). Response Conflict as an Environmental Determinant of Gender Differences in Sensitivity to Alcohol Impairment. Current Drug Abuse Reviewse, 3(3), 147–155. 10.2174/1874473711003030147

Wiers, C. E., Stelzel, C., Park, S. Q., Gawron, C. K., Ludwig, V. U., Gutwinski, S., Heinz, A., Lindenmeyer, J., Wiers, R. W., Walter, H., & Bermpohl, F. (2014). Neural Correlates of Alcohol-Approach Bias in Alcohol Addiction: the Spirit is Willing but the Flesh is Weak for Spirits. Neuropsychopharmacology, 39(3), 688–697. 10.1038/npp.2013.252

Wiers, R. W., Eberl, C., Rinck, M., Becker, E. S., & Lindenmeyer, J. (2011). Retraining Automatic Action Tendencies Changes Alcoholic Patients’ Approach Bias for Alcohol and Improves Treatment Outcome. Psychological Science, 22(4), 490–497. 10.1177/0956797611400615

Wobbrock, J. O., Findlater, L., Gergle, D., & Higgins, J. J. (2011). The Aligned Rank Transform for nonparametric factorial analyses using only ANOVA procedures. Conference on Human Factors in Computing Systems - Proceedings, 143–146. 10.1145/1978942.1978963

Xia, X., Li, Y., Wang, Y., Xia, J., Lin, Y., Zhang, X., Liu, Y., & Zhang, J. (2021). Functional role of dorsolateral prefrontal cortex in the modulation of cognitive bias. Psychophysiology, 58(10). 10.1111/psyp.13894

Yamada, K., & Toda, K. (2023). Habit formation viewed as structural change in the behavioral network. Communications Biology, 6(1), 303. 10.1038/s42003-023-04500-2

Zhang, G., Garrett, D. R., & Luck, S. J. (2024). Optimal filters for ERP research II: Recommended settings for seven common ERP components. Psychophysiology, 61(6), e14530. 10.1111/PSYP.14530

