## Supplementary Material for "Modulation of Automatic Alcohol Approach Tendencies using Single-Session 10 Hz rTMS over the Right dLPFC"

### Exploratory Analysis of Participants with Baseline Alcohol Avoidance Tendencies

Given the limited sample size of participants exhibiting baseline alcohol avoidance tendencies (AAI = −0.057 ± 0.021; n = 9), all such individuals were assigned to the active rTMS condition, with no sham comparison for this group. Accordingly, the analyses presented below are exploratory and aim to characterize (i) baseline neural responses and (ii) behavioral and neural changes following stimulation within this subgroup.

#### Baseline Characteristics

In contrast to the approach group, which showed robust alcohol-specific attenuation in both RF N2 and CP P3b amplitudes, the avoidance group exhibited a more restricted pattern of stimulus differentiation at baseline. Specifically, RF N2 amplitudes were significantly reduced for alcohol relative to non-alcohol cues (median difference = 2.04, $W = 259$, $p = 0.035$, $r = 0.416$), suggesting attenuated prefrontal cognitive control in response to alcohol stimuli. However, this effect was not observed at other sites, i.e., neither LF N2 (median difference = 0.49, $W = 133$, $p = 0.286$, $BF_{10} = 0.375$) nor CP P3b amplitudes (median difference = −1.82, $W = 105$, $p = 0.075$, $BF_{10} = 0.787$) showed significant stimulus-related differences. Overall, these findings indicate that alcohol-specific neural differentiation in the avoidance group was limited to prefrontal regions and did not extend to broader attentional processing systems, as observed in the approach group.

Importantly, no stimulus × action interaction was observed at any region (LF: $F(1, 8) = 0.763$, $p = 0.408$, $BF_{10} = 0.068$; RF: $F(1, 24) = 1.165$, $p = 0.312$, $BF_{10} = 0.099$; CP: $F(1, 8) = 4.665$, $p = 0.063$, $BF_{10} = 0.217$), indicating that baseline neural responses to alcohol cues did not vary as a function of action type.

#### Post-rTMS Behavioral Effects

Following active 10 Hz rTMS, the avoidance group showed a significant increase in AAI (median difference = 0.126, $W = 43$, $p = 0.012$, $r = 0.593$), reflecting a shift from avoidance toward approach tendencies (Figure 1). This pattern contrasts with the approach group, in which rTMS reduced AAI. Notably, these opposing effects resulted in convergence of AAI scores between groups post-rTMS ($W = 64$, $p = 0.525$, $BF_{10} = 0.48$), suggesting that the direction of rTMS-induced behavioral change depends on baseline motivational orientation. To identify the behavioral components underlying this shift, response times were examined across the four trial types (alcohol pull, alcohol push, non-alcohol pull, non-alcohol push). No significant pre–post changes were observed in any individual trial type (all $p > 0.05$; Figure 2, Panels A and B). This indicates that the increase in AAI was not driven by a specific component but rather reflects distributed, small changes across multiple trial types.

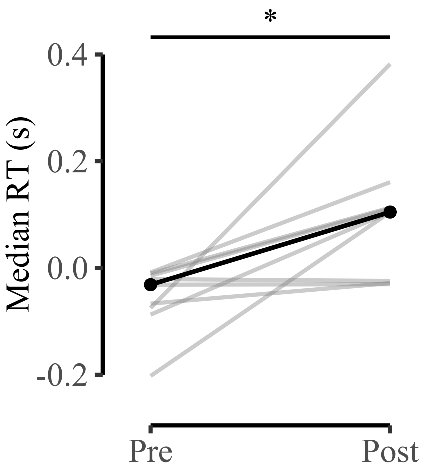


***Figure 1****. Active 10 Hz stimulation increased AAI in individuals with baseline avoidance tendencies, reflecting a shift from avoidance to approach tendencies. Lighter lines represent individual data, and the darker line indicates the group median.*

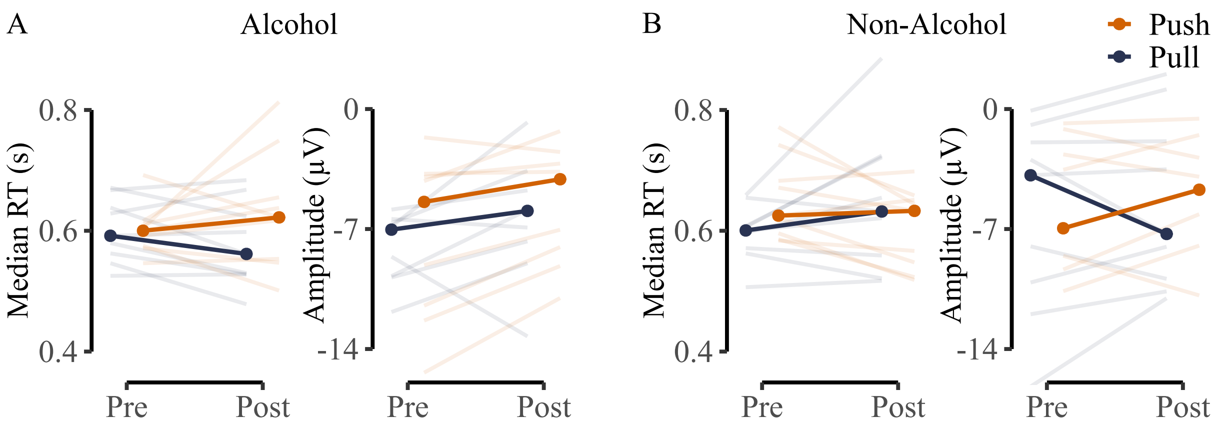


***Figure 2****. No action- or stimulation-specific modulation was observed in response times (RTs) and RF N2 amplitudes in the avoidance group for alcohol (Panel A) and non-alcohol stimuli (Panel B). Lighter lines represent individual participants, and darker lines indicate medians.*

**Post-rTMS Neural Effects**

Within the avoidance group, despite baseline attenuation of RF N2 amplitudes for alcohol stimuli, rTMS did not significantly modulate RF N2 as a function of stimulus type. No significant stimulus (alcohol vs non-alcohol) × assessment phase (pre vs post stimulation) interaction ($F(1, 56) = 1.520$, $p = 0.223$, $BF_{10} = 0.492$), or stimulus × action × assessment phase interaction ($F(1, 56) = 0.152$, $p = 0.698$, $BF_{10} = 0.397$; Figure 2 Panels A and B) was observed. These findings indicate that post-rTMS changes in prefrontal cognitive control did not differ between alcohol and non-alcohol cues across pull and push actions. Thus, although the avoidance group exhibited alcohol-specific RF N2 attenuation at baseline, this pattern was not further modulated by 10 Hz rTMS targeting the right dLPFC.
